# Dynamic DNA-based information storage

**DOI:** 10.1101/836429

**Authors:** Kevin N. Lin, Albert J. Keung, James M. Tuck

## Abstract

Technological leaps are often driven by key innovations that transform the underlying architectures of systems. Current DNA storage systems largely rely on polymerase chain reaction, which broadly informs how information is encoded, databases are organized, and files are accessed. Here we show that a hybrid ‘toehold’ DNA structure can unlock a fundamentally different, dynamic DNA-based information storage system architecture with broad advantages. This innovation increases theoretical storage densities and capacities by eliminating non-specific DNA-DNA interactions common in PCR and increasing the encodable sequence space. It also provides a physical handle with which to implement a range of in-storage file operations. Finally, it reads files non-destructively by harnessing the natural role of transcription in accessing information from DNA. This simple but powerful toehold structure lays the foundation for an information storage architecture with versatile capabilities.

## Introduction

The creation of digital information is rapidly outpacing conventional storage technologies^1^. DNA may provide a timely technological leap due to its high storage density, longevity^2–4^, and energy efficiency^5^. A generic DNA-based information storage system is shown in Figure 1A, where digital information is encoded into a series of DNA sequences, synthesized as a pool of DNA strands, read by DNA sequencing, and decoded back into an electronically compatible form. A growing body of work has focused on implementing and improving each of these four steps^6–11^; however, a relative dearth of research has explored technologies to dynamically access and manipulate data within storage databases^12,13^. This is likely because DNA synthesis and sequencing are considerably slower processes than electronic writing and reading of data^14^; thus, DNA would likely serve at the level of archival or cold storage where information could be stored and infrequently accessed from a relatively static DNA database. Yet, an archival DNA database, just like electronic versions, need not be completely static and would benefit greatly from dynamic properties^15,16^. For example, file operations could facilitate in-storage computation, while the ability to non-destructively and repeatedly access the database would reduce DNA synthesis costs and abrogate the need to store multiple copies of archives. If possible, implementation of dynamic properties would bring DNA-based storage systems one step closer to practical viability.

**Figure 1:**
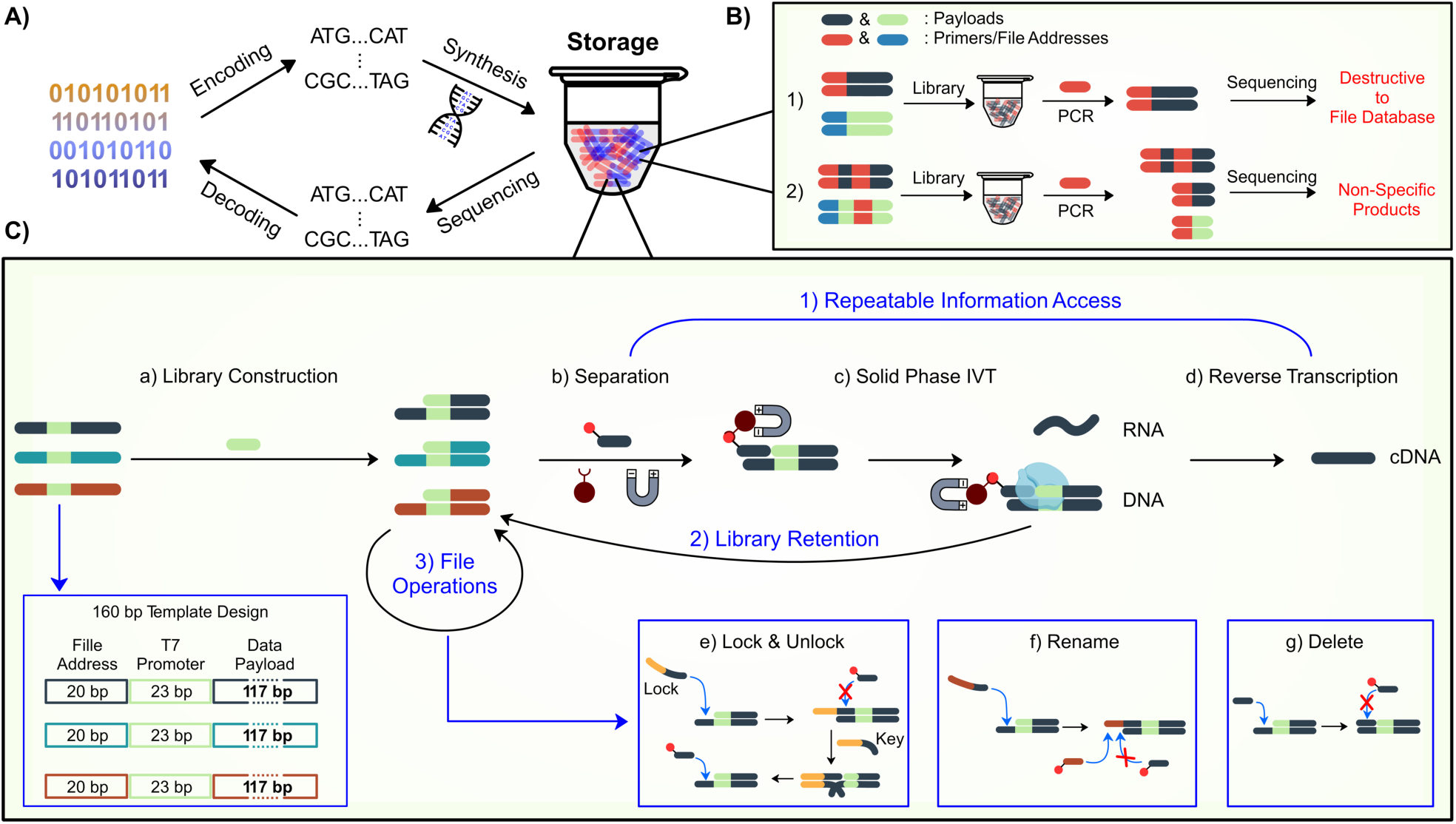
Molecular technologies unlock dynamic operations for DNA storage. **A)** The generic framework for DNA-based storage systems includes encoding of digital information to nucleotide sequences, DNA synthesis and storage, DNA sequencing, and decoding the desired information. **B)** Schematic of challenges faced by PCR-based file access. **C)** A schematic of DORIS (Dynamic Operations and Reusable Information Storage). A toehold-based hybrid DNA structure enables repeatable information access through non-PCR based magnetic separation, *in-vitro* transcription, reverse transcription, and the return of separated files to the database. Additionally, toeholds unlock in-storage file operations including ‘lock’, ‘unlock’, ‘rename’, and ‘delete’.

A key requirement but major challenge for engineering dynamic properties into storage systems is that they would require methods that do not cause undesired permanent alterations or loss of data. Thus, we took inspiration from how biological systems access genomic information, as well as from the fields of DNA computing and synthetic biology. In this work, we describe a dynamic DNA-based storage system that harnesses technologies from molecular biology to implement reusable and repeatable file access and file operations including file locking, renaming, and deletion. In addition, as positive byproducts, this dynamic system improves file access specificity and reduces data encoding overhead, thus increasing information density and theoretical maximum capacity. This work demonstrates dynamic information access and manipulations are practical for DNA-based information storage systems. For convenience, we refer to this system collectively as DORIS (Dynamic Operations and Reusable Information Storage).

## Result

### Strategy: molecular technologies to unlock dynamic features for DNA storage

One of the fundamental “dynamic” features of information storage systems is the ability to access specific data or files from a database. Current methods in DNA-based storage systems rely on PCR where primers are used to copy the DNA strands of a desired file. However, PCR presents three challenges that may preclude its broader use as a platform technology (Figure 1B): 1) PCR amplifies and enriches specific DNA strands for downstream DNA sequencing, and either destroys the entire database or significantly alters its composition; 2) Since dsDNA templates are melted in each PCR cycle, primers specific for a desired file may bind non-specifically to similar sequences in any region of any strand in the database. This can lead to undesired off-target products being generated and requires encodings that sacrifice information density in order to avoid conflicting DNA sequences appearing in DNA strands^7^. In fact, our simulations find that this limitation restricts theoretical densities and capacities by over 30 fold when dense encodings are used (Figure 2D), with some densities not even achievable without encountering primer conflicts within the data payload region; 3) Finally, it is unclear how PCR, a method that actively creates new DNA strands, can serve as the foundation for in-storage file manipulations without significant challenges associated with altering the composition and strand balance of a database. To address this set of challenges, we propose a simple but fundamental shift in DNA storage system architectures, inspired by single stranded DNA (ssDNA) ‘toeholds’ used in synthetic biology and DNA computing^17–20^, and by the way organisms naturally access information from their genome through transcription. As described below and in Figure 1C, we engineered a reusable DNA-based information storage system that can be created at scale, reduces off-target file access, and supports multiple in-storage file operations.

**Figure 2.**
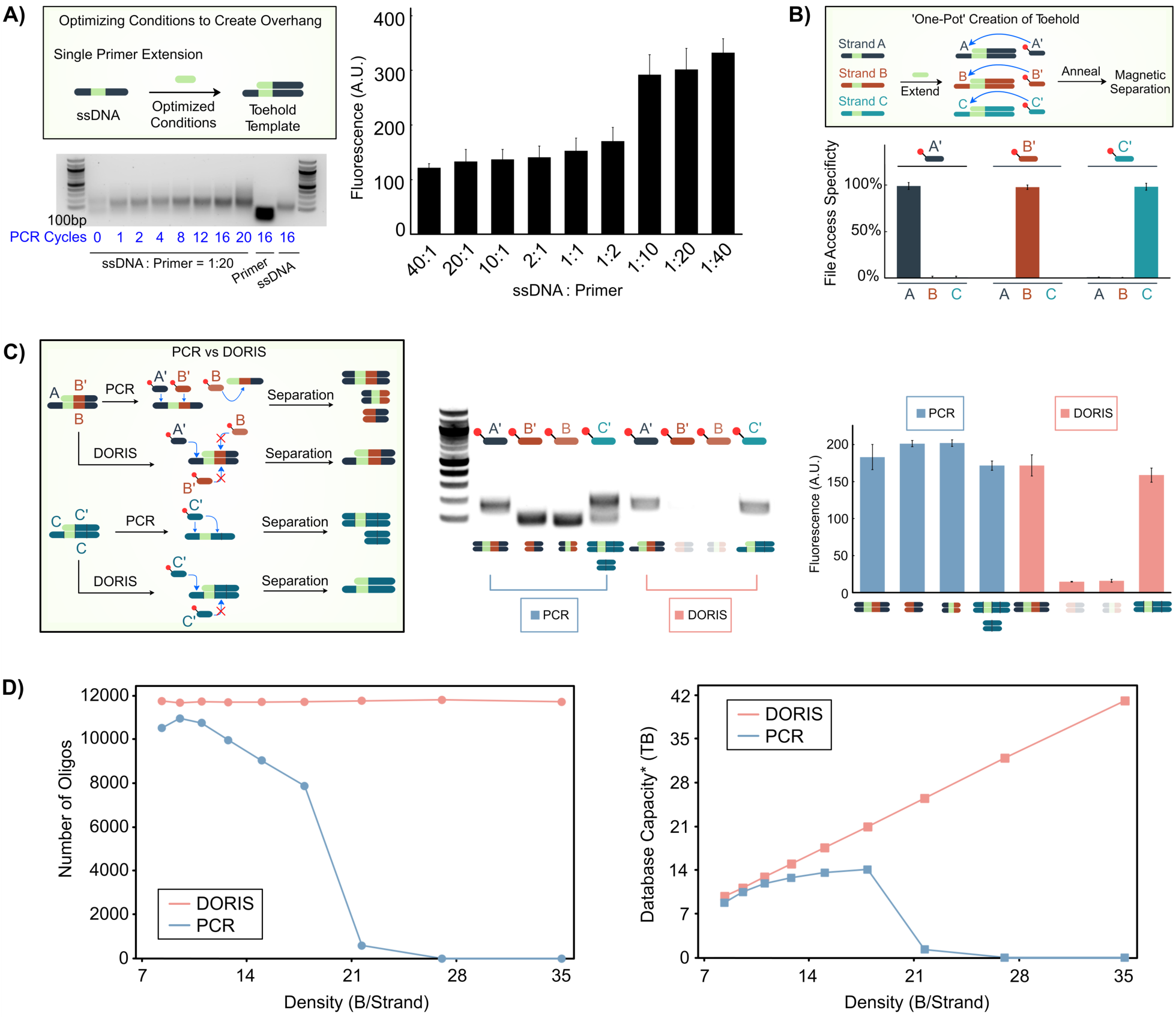
Toehold-based file access eliminates non-specific interactions and increases density and capacity limits. **A)** Single primer extension creates hybrid DNA toehold strands. **(Bottom)** 4 cycles of PCR generated the optimal amount of 160 bp hybrid toehold strands while minimizing excess ssDNA production. **(Right)** DNA gel fluorescence showed a marked increase in generation of toeholds below 1:10 ssDNA:primer ratios. **B)** Individual files can be accessed from a 3-file database created by a ‘one-pot’ single primer extension. Each file was accessed by its corresponding biotin-linked oligo, followed by a non-PCR based separation using functionalized magnetic beads. File access specificity is the percentage of the DNA accessed by a biotin-linked oligo that is either file A, B, or C as measured by qPCR. **C) (Left)** PCR but not DORIS will allow file-access oligos to bind internal off-target sites and produce undesired products. **(Middle)** DNA gels and (**Right)** their quantified fluorescence show that PCR-based access results in truncated and undesired amplicons whereas DORIS accesses only the desired strands. **D) (Left)** Monte Carlo simulations estimated the number of oligos found that will not interact with each other or the data payload. 20,000 oligos were tested for different density encodings. The x-axis represents density, which is inversely related to the length of ‘codewords’ used to store discrete 1 byte data values. We evaluated codeword lengths of 12 through 4. For a PCR-based system, longer codewords have lower density but less diverse overall data payload sequences, thus allowing more distinct oligos to be used as addresses in a database. For DORIS, the encoding density is not impacted because it need not guard against undesired binding between the oligo and data payloads. **(Right)** For PCR, the number of oligos that will not bind the data payload drops as strand density increases, which means that fewer files can be stored, leading to a lower overall system capacity. For DORIS, the availability of oligos is independent of encoding, and capacity therefore increases with denser encodings. Error bars are standard deviations of three replicate file accesses. *Capacities may be limited by synthesis and sequencing limitations not accounted for here.

### Toehold structures can be efficiently created in ‘one-pot’ reactions

The toehold serves as the central architectural feature of the system: DNA strands are comprised of dsDNA with 3’ ssDNA overhangs (Figure 1C). We envisioned that toeholds could be multipurpose structures serving as file addresses as well as molecular ‘handles’ for file operations. As future DNA databases would be comprised of upwards of 10^15^ distinct strands^21^, we first asked if toeholds could be created in a high throughput and parallelized manner. We began with 160 bp ssDNA strands with a common 23 bp sequence inset 20 bp from the 3’ end (Figure 1C, Table S1, Figure 2A). This 23 bp sequence contains the T7 RNA polymerase binding sequence (allowing data access directly by T7-based *in vitro* transcription, described below). Here we used the sequence to bind a primer that would be common to all strands in a database, followed by several cycles of thermal annealing and DNA polymerase extension (e.g. ‘PCR cycles’ but with only one primer), resulting in toehold strands with a 20 bp 3’ ssDNA overhang (Figure 2A, top). We optimized the ratio of ssDNA to primer, the number of cycles, along with other system parameters (Figure S1) to maximize the amount of ssDNA converted to toehold structures. We found that decreasing the ssDNA:primer ratio past 1:10 led to a step change in the amount of toeholds produced as quantified by gel electrophoresis (Figure 2A). We decided to conservatively work with a 1:20 ssDNA:primer ratio; at that ratio we found that only 4 PCR cycles were needed to convert the ssDNA into toehold structures, as seen by the upward shift in a DNA gel (Figure 2A).

Next, we tested whether this method could be used to create 3 distinct toehold strands in ‘one pot’ and if each strand could then be specifically separated from the mixture (Figure 2B). We mixed 3 distinct ssDNAs “A”, “B”, and “C” together, added T7 primer, and performed 4 PCR cycles. We then used biotin-linked, 20 bp DNA oligos to bind each strand (i.e. “file”) and separated them out from the mixture using streptavidin-linked magnetic beads. Each of these “file-access” oligos were able to specifically pull out only their corresponding strands without the other two (Figure 2B, bottom). Importantly, this file separation step did not require repetitive and high temperature annealing steps, and was performed at room temperature (25 °C) with only minimal gains observed at higher oligo annealing temperatures of 35 or 45 °C (Figure S2).

While 20 bp is a standard DNA primer or oligo length, we asked what effects toehold length and access temperature would have on file access efficiency. We designed 5 strands with 5-25 bp toeholds (Figure S3). We then accessed each strand using its specific biotin-linked oligo and magnetic separation at 15-55 °C. We observed an enhanced access efficiency for longer toehold sequences (20bp and 25bp) and at lower temperatures (Figure S3B). This was in agreement with a thermodynamic analysis (Figure S3C).

### Toehold-based file access eliminates non-specific interactions and increases density and capacity limits

We hypothesized that the direct access of files through toehold structures could provide an advantage over PCR-based file access: by eliminating the need to thermally anneal the system, the strands would not denature and could act as a natural barrier to oligos binding non-specifically within data payload regions. This would increase the theoretical information density and capacity of storage systems by allowing sequences similar to the oligos (file addresses) to appear in data payloads. To compare DORIS with PCR-based access, we created two toehold strands (Figure 2C). One strand had an internal binding site for oligo B’ and a toehold that bound oligo A’. We hypothesized and experimentally verified that by using DORIS, only oligo A’ but not oligo B’ could access the strand (Figure 2C). In contrast, when PCR was used, both oligo A’ and oligo B’ accessed the strand, with oligo B’ producing undesired truncated products. The second strand we tested had an internal binding site and toehold that both bound oligo C’. We showed that using DORIS with oligo C’ separated out the full-length strand. In contrast, when using PCR, oligo C’ created both full length and truncated strands.

To assess the impact of the increased file access specificity DORIS provides, we performed Monte Carlo simulations to estimate the total number of oligo sequences and total capacities achievable when oligo sequences were or were not prohibited from appearing in the data payload regions (Figure 2D). The data payload region can be made more or less diverse in sequence, which corresponds to more or less information density, respectively. However, with increased diversity and density, the likelihood of an oligo sequence appearing in the data payload increases; for PCR (but not DORIS) this reduces the number of primers available to be used, leading to a reduction in total system capacity. We found that capacity monotonically increases with increasing density for DORIS; in contrast, for PCR, increasing density initially provides a minor benefit to overall capacity, but eventually leads to a catastrophic drop in capacity as the number of non-conflicting primers quickly drops to zero. It is possible to increase the number of strands per file as encoding density is increased to make up for the loss of primers. However, this runs counter to the goals of random access since it would result in files too large to sequence and decode in a single sequencing run. It is important to note that our simulations are based upon very conservative database sizes of only 10^9^ DNA strands, while future storage systems are likely to exceed 10^12^ strands or greater. As database sizes and the total amount of DNA sequence space increase, the number of primers available for PCR-based systems will drop even further while DORIS will be unaffected. Therefore, the theoretical capacity and density improvements DORIS provides could be orders of magnitude greater than what is estimated in our simulations. It should also be noted that we have artificially constrained the total number of codewords in each system to be limited to 256 due to practical considerations for the simulation. For longer codewords, there are many more theoretically possible codewords (4^codeword length^), therefore this constraint of 256 codewords substantially reduces the total capacity of the system. We anticipate capacity differences between DORIS and PCR to be substantially greater at longer codeword lengths than what we report in Figure 2D if more codewords are allowed. In summary, a database comprised of toehold structures can be created efficiently in ‘one pot’, and toeholds facilitate a non-PCR based file access method that enhances file access specificity and increases theoretical database densities and capacities.

### DORIS mimics natural transcription to non-destructively access information

The next challenge we addressed was how to access a file’s information without needing to destroy the file itself through the DNA sequencing process. It would be preferable to return the file to the database for future use. For inspiration, we looked to natural biological systems where information is repeatedly accessed from a single permanent copy of genomic DNA through the process of transcription. The T7 promoter we had designed to be common to all strands served not only the purpose of synthesizing the toehold structures but also of initiating transcription^22^. Our strategy was to physically retrieve a file of interest using biotin-linked oligos and streptavidin coated magnetic beads, *in vitro* transcribe (IVT) RNA directly from the bead-coupled file, return the file to the library, and reverse transcribe the RNA into cDNA for downstream analysis or sequencing (Figure 3A).

**Figure 3:**
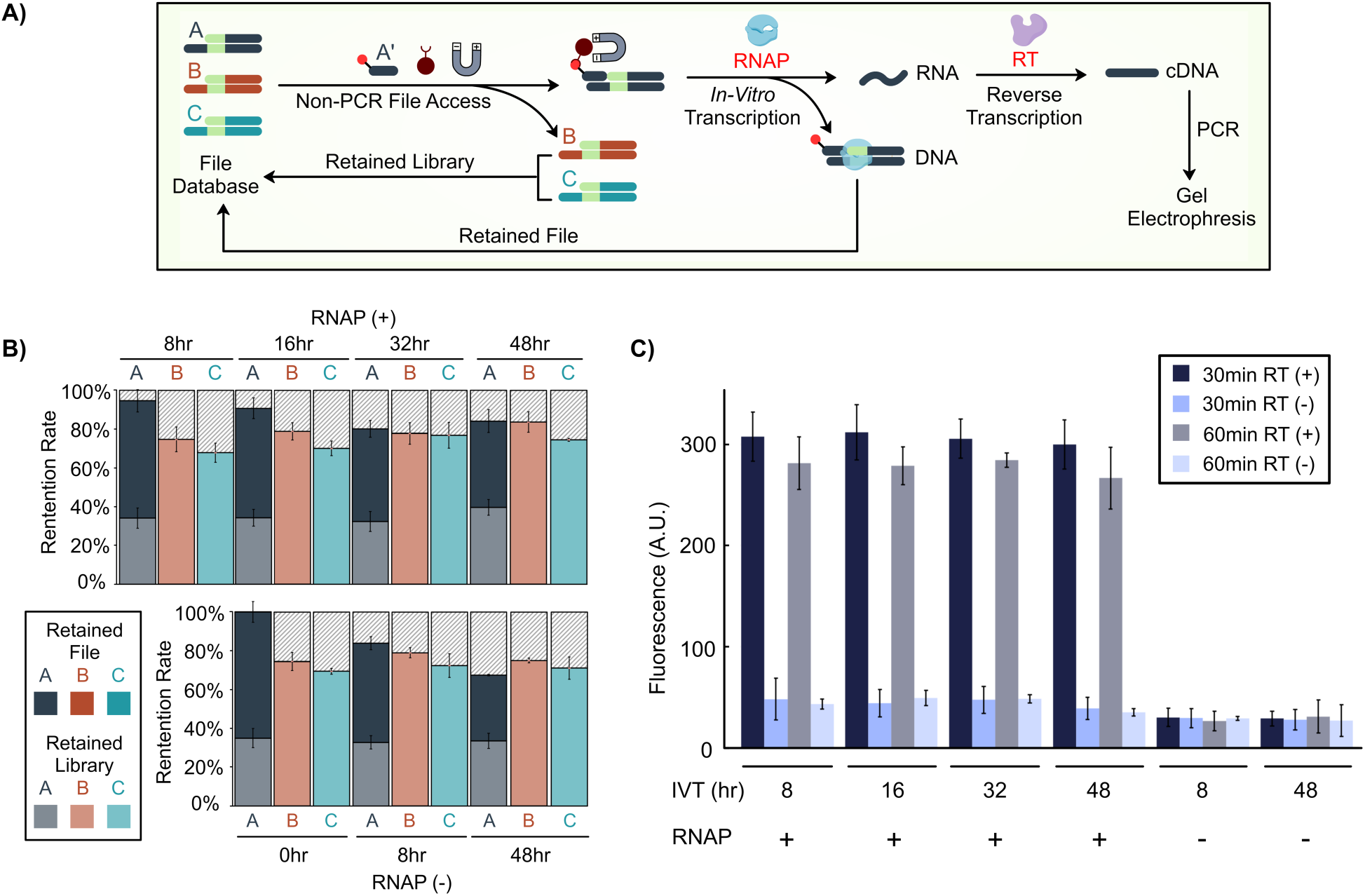
DORIS mimics natural transcription to non-destructively access information. **A)** File A was separated using non-PCR based magnetic separation while the library was recovered (‘Retained Library’). T7-based *in vitro* transcription was performed directly on the bead-immobilized file for up to 48 hours to generate RNA. Reverse transcription converted the RNA to complementary DNA (cDNA) while the immobilized file A was released back into the database (‘Retained File’). **B)** The amount of Retained Library and Retained File (after file A was accessed by oligo A’) was measured by qPCR and plotted as a percentage of the original amount of each file that was in the database. The specificity of file access is evident by the absence of file B and C in the Retained File. The presence of T7 RNA polymerase (RNAP) did not affect the retention of file A. **C)** cDNA generated from accessed file A was amplified by PCR to sub-saturating quantities, run on a DNA gel, and quantified by SYBR green fluorescence. RT (reverse transcriptase). IVT (*in vitro* transcription). Error bars are standard deviations of three replicate file accesses.

We implemented this system with 3 distinct toehold strands representing a file database and accessed file A with a biotin-linked oligo A’. We then measured the amounts and compositions of the “retained library” and “retained file” (Figure 3B). The retained library had higher levels of files B and C compared to A, as some of the file A strands had been magnetically removed. The retained file contained only file A strands, with no B or C. The net total amount of file A recovered from the retained library and retained file was 95% of what was originally in the database. This suggests that the loss of file A throughout the process was minimal. We asked if the length of time of the IVT would affect the amount of recovered DNA, as IVT operates at an elevated temperature of 37 °C and could destabilize DNA. There was a slight trend of DNA loss with longer incubation times of up to 48 hours; however, this loss could be improved by adding an annealing step after IVT (Figure S4). Furthermore, we checked that the information from file A could be retrieved following IVT by reverse transcription and PCR amplification: we found that increasing IVT times from 8 to 48 hours improved the yield of RNA (Figure S5) but the reverse transcription step was likely saturated as it did not improve the yield of dsDNA subsequently created (Figure 3C); therefore, 8-hour IVTs are sufficient for maximum file access. We also found that 60-minute reverse transcriptions had no beneficial effects over 30 minutes. Finally, leaving out either T7 RNA polymerase or reverse transcriptase abrogated file access, indicating that the information recovered was specifically derived from RNA production and not due to any contaminating strands from the original database (Figure 3C). Overall, DORIS is able to access specific information while retaining significant amounts of the original database and accessed file.

### Toeholds enable in-storage file operations

Many inorganic information storage systems, even cold storage archives, maintain the ability to dynamically manipulate files. Similar capabilities in DNA-based systems would significantly increase their value and competitiveness. DNA has previously been used to execute computations^23–25^, and we therefore hypothesized toeholds could be used to implement in-storage file operations. As a proof-of-principle, we implemented locking, unlocking, renaming, and deleting files and showed these operations could be performed at room temperature (Figure 4).

**Figure 4:**
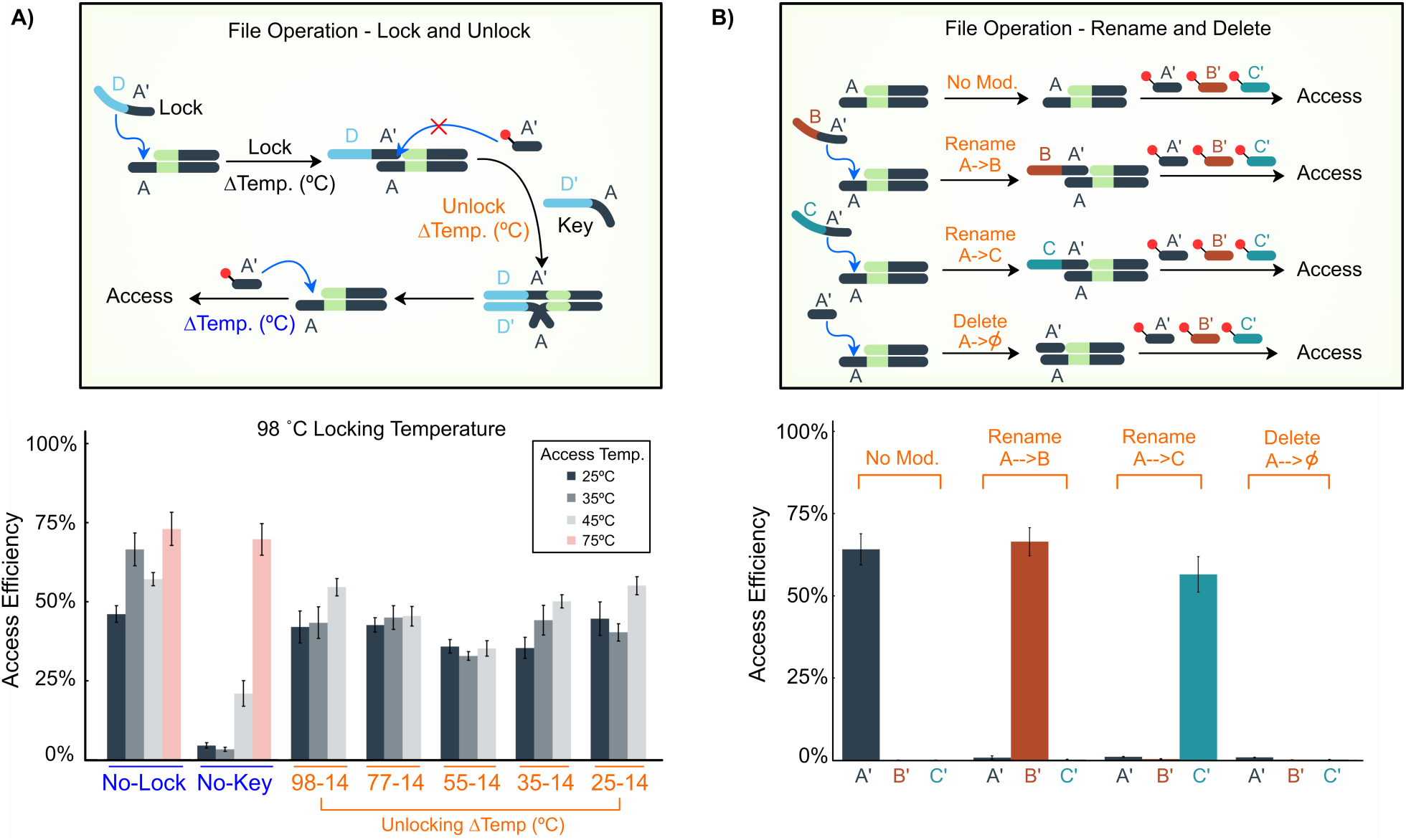
Toeholds enable in-storage file operations. **A) (Top)** Schematic of ‘locking’ and ‘unlocking’ in-storage file operations. **(Bottom)** Attempts to access file A by DORIS without locking (‘No-Lock’), with locking but without a key (‘No-Key’)., or with locking and key added at different temperatures (orange). The lock was added at 98 °C. The key was added at different temperatures (orange) and then cooled to 14 °C. The accessing oligo A’ was added at different access temperatures of 25, 35, 45, or 75 °C for 2 minutes, followed by a temperature drop of 1 °C/minute to 25 °C. Access efficiency is the amount of file A recovered relative to its original quantity, as measured by qPCR. **B) (Top)** Schematic of ‘rename’ and ‘delete’ operations. File A was modified by renaming or deleting oligos. **(Bottom)** The completion of each operation was tested by measuring how much of the file was access by each separate oligo: A’, B’, or C’. Access efficiency is the amount of file A accessed relative to its original amount, as measured by qPCR. ‘No Mod’ (No file modification/operation). Error bars are standard deviations of three replicate file operations/accesses.

We started with the 3-file database and tested the ability of a biotin-linked oligo A’ to bind and access file A at a range of temperatures from 25 to 75 °C (Figure 4A, bottom, ‘no lock’). File A was successfully accessed at all temperatures with roughly 50% of its strands accessed. To lock file A, we extracted file A from the 3-file database and mixed in a long 50 bp ssDNA ‘lock’ that had a 20bp complementary sequence to the toehold of file A. With the lock in place, oligo A’ was no longer able to access the file except at higher temperatures above 45 °C (‘no-key’), presumably because the lock was melted from the toehold, allowing for oligo A’ to compete for toehold binding. To unlock the file, we added the ‘key’ that was a 50 bp ssDNA fully complementary to the lock. We tested different unlocking temperatures and found the key was able to remove the lock at room temperature with the same efficiency as at higher temperatures. This is likely due to the long 30 bp toehold presented by the lock, allowing the key to ‘unzip’ the lock from file A. We also optimized the relative molar ratios (file A:lock:key:oligo A’ = 1:10:10:15) to minimize off-target access and ensure proper locking. We did observe that the temperature at which the lock was added influenced the fidelity of the locking process. At 98 °C, the locking process worked well. When the lock was added at 25 °C, there was leaky access even when no key was added (Figure S6). This may be due to secondary structures preventing some file A strands from hybridizing with locks at low temperatures. Fortunately, locking at 45 °C had reasonable performance, thus avoiding the need to elevate the system to 95 °C. In the context of a future DNA storage system, files could first be extracted then locked or renamed at an elevated temperature, then returned to the database, thus avoiding exposure of the entire database to elevated temperatures. The entire process could otherwise be performed at room temperature.

We also implemented file renaming and deletion. To rename a file with address A to have address B, we mixed file A with a 40 bp ssDNA that binds to A, with the resultant toehold being address B (Figure 4B). We added all components at similar ratios to the locking process (file:renaming oligo:accessing oligo = 1:10:15) and the renaming oligo was added at 45 °C. We then tested how many file strands each oligo A’, B’, or C’ could access and found that the renaming process completely blocked oligos A’ or C’ from accessing the file (Figure 4B, bottom). Only oligo B’ was able to access the file suggesting that almost all strands were successfully renamed from A to B. Similarly, we successfully renamed file A to C. Based on the ability of renaming oligos to rename files with near 100% completion, we hypothesized and indeed found that a short 20 bp oligo fully complementary to A could be used to completely block the toehold of file A and essentially ‘delete’ it from the database (Figure 4B, bottom). A file could also simply be extracted from a database to delete it as well. However, this alternative form of ‘blocking’ deletion suggests one way to ensure any leftover file strands that were not completely extracted would not be spuriously accessed in the future.

## Discussion

DORIS represents a proof of principle framework for how inclusion of a few simple innovations can fundamentally shift the physical and encoding architectures of a system. In this case, the toehold structure drives multiple new capabilities for DNA storage: 1) it increases the theoretical information density and capacity of DNA storage by inhibiting non-specific binding within data payloads; 2) it enables transcription-based, non-destructive information access; and 3) it makes possible in-storage file operations. We envision other new architectures and capabilities may be on the horizon given rapid advances in DNA origami^26^, molecular handles^27–29^, and molecular manipulations developed in fields such as synthetic biology^30^.

Beyond the specific capabilities enabled by DORIS, one of the greatest benefits we envision DORIS providing is compatibility with future miniaturized and automated devices^31,32^. In particular, DORIS can operate with no temperature cycling and functions at or close to room temperature for all steps. This has potential advantages for maintaining DNA integrity and database stability while also simplifying the design of future automated DNA storage system devices. In addition, a single DNA database sample can be ‘reused’ rather than having to store many physical copies that are destroyed upon each file query. It is also intriguing to consider how in-storage operations like ‘lock’ & ‘unlock’ are not common “hardware” operations in conventional storage technologies, and may therefore offer unique unforeseen capabilities in the future. All of these features lend DORIS to be easily translated to systems with automated fluid handling and magnetic actuation.

DORIS is also a fundamentally scalable system. The creation of toehold strands is simple and high throughput, it is compatible with existing file system architectures including hierarchical addresses^11,21^, and it facilitates scaling of capacity. While the need to include the T7 promoter in every strand does occupy valuable data payload space, it is a worthwhile tradeoff; while the T7 promoter decreases data density and capacity in a linear fashion, it improves both metrics exponentially by allowing many sequences to appear in the data payload that normally would have to be avoided in PCR-based systems (or conversely by allowing the full set of mutually ‘good’, non-conflicting primers to be used)^11,21^. Future work may assess how DORIS and other physical innovations may alter and reduce the stringency of encoding and error correction algorithms and subsequently benefit system density and capacity.

Of course, as with all information storage systems, there are challenges and questions regarding the efficiency and accuracy of each technology that will be important to address prior to commercial implementation. For example, future work might assess how each step of DORIS performs in the context of a highly diverse and dense database of strands, both in terms of efficiency and informational retrieval error rates. In particular, new materials, evolved enzymes, and the optimization of reaction conditions could improve DNA recovery percentages to drive DORIS towards a fully reusable system. Currently, recovery rates are between 50-100% (Figure 3B), meaning multiple copies of each strand would need to be included in the system, and that each physical database would eventually need to be replenished after a finite number of file queries. Devoting resources and attention to such optimizations need to be balanced with the fact that the field of molecular information storage is nascent and that there are likely a wide range of new capabilities and physical innovations that could be explored and introduced into the field.

Finally, we believe this work, through the concept of in-storage computation, motivates a merging of work in the fields of DNA computation, synthetic biology, and DNA storage. In-storage computation and file operations could increase the application space of DNA storage, or identify novel applications areas, such as in the highly parallel processing of extreme levels of information (e.g. medical, genomic, and financial data). DORIS complements and harnesses the benefits of prior work while providing a feasible path towards future systems with advanced, hybrid capabilities.

## Methods

### Creation of toeholds strands

Toehold strands were created by ‘filling in’ ssDNA templates (IDT) with primer TCTGCTCTGCACTCGTAATAC (Eton Bioscience) at a ratio of 1:40 using 0.5 µL of Q5 High-Fidelity DNA Polymerase (NEB, M0491S) in a 50 µL reaction containing 1x Q5 polymerase reaction buffer (NEB, B9072S) and 2.5 mM each of dATP (NEB, N0440S), dCTP (NEB, N0441S), dGTP (NEB, N0442S), dTTP (NEB, N0443S). The reaction conditions were 98 °C for 30 s and then 4 cycles of: 98 °C for 10 s, 53 °C for 20 s, 72 °C for 10 s, with a final 72 °C extension step for 2 min. Toehold strands were purified using AMPure XP beads (Beckman Coulter, A63881) and eluted in 20 μL of water.

### File separations

Oligos were purchased with a 5’ biotin modification (Eton Bioscience). Toehold strands were diluted to 10^11^ strands and mixed with biotinylated oligos at a ratio of 1:40 in a 50 µL reaction containing 2 mM MgCl_2_ (Invitrogen, Y02016) and 50 mM KCl (NEB, M0491S). Oligo annealing conditions were 45 °C for 2 min, followed by a temperature drop at 1 °C/min to 14 °C. Streptavidin magnetic beads (NEB, S1420S) were prewashed using high salt buffer containing 20 mM Tris-HCl, 2 M NaCl and 2 mM EDTA pH 8 and incubated with toehold strands at room temperature for 30 min. The retained library was recovered by collecting the supernatant of the separation. The beads were washed with 100 µL of high salt buffer and used directly in the *in vitro* transcription reaction. After transcription, the beads with the bound files were washed twice with 100 µL of low salt buffer containing 20 mM Tris-HCl, 0.15 M NaCl and 2 mM EDTA pH 8 and subsequently eluted with 95% formamide (Sigma, F9037) in water. The quality and quantity of the DNA in the retained library and file were measured by quantitative real time PCR (Bio-Rad).

### *In vitro* transcription

Immobilized toeholds strands bound on the magnetic beads were mixed with 30 µL of *in vitro* transcription buffer (NEB, E2050) containing 2 µL of T7 RNA Polymerase Mix and ATP, TTP, CTP, GTP, each at 6.6 mM. The mixture was incubated at 37°C for 8, 16, 32, and 48 hours, followed by a reannealing process where the temperature was reduced to 14 °C at 1 °C/min to enhance the retention of toeholds on the beads. The newly generated RNA transcripts were separated from the streptavidin magnetic beads and their quantity measured using the Qubit RNA HS Assay Kit (Thermo Fisher, Q32852).

### Reverse transcription

First-strand synthesis was generated by mixing 5 µL of separated RNA transcript with 500 nM of reverse primer in a 20 µL reverse transcription reaction (Bio-Rad, 1708897) containing 4 µL of reaction supermix, 2 µL of GSP enhancer solution and 1 µL of reverse transcriptase. The mixture was incubated at 42 °C for 30 or 60 mins, followed by a deactivation of the reverse transcriptase at 85 °C for 5 min. The resultant cDNA was diluted 100-fold, and 1 µL was used as the template in a PCR amplification containing 0.5 µL of Q5 High-Fidelity DNA Polymerase (NEB, M0491S), 1x Q5 polymerase reaction buffer (NEB, B9072S), 0.5uM of forward and reverse primer, 2.5 mM each of dATP (NEB, N0440S), dCTP (NEB, N0441S), dGTP (NEB, N0442S), dTTP (NEB, N0443S) in a 50 µL total reaction volume. The amplification conditions were 98 °C for 30 s and then 25 cycles of: 98 °C for 10 s, 55 °C for 20 s, 72 °C for 10 s with a final 72 °C extension step for 2 min. The quality of amplification was measured using gel electrophoresis.

### Locking and unlocking

Lock and key strands were purchased from Eton Biosciences. To lock the file, purified toehold strands were mixed with lock strands at a molar ratio of 1:10 in a 25 µL reaction containing 2 mM MgCl_2_ and 50 mM KCl. The mixture was annealed to 98 °C, 45 °C or 25 °C for 2 min, followed by a temperature drop at 1 °C/min to 14 °C. To unlock the file, key strands were added into the locked file mixture at a molar ratio of 10:1 to the original toehold strand amount. The mixtures were annealed to 98, 77, 55, 35, or 25 °C for 2 min, followed by a temperature drop at 1 °C/min to 14 °C. To access the unlocked strands, file-specific biotin-modified oligos were added into the mixture at a ratio of 15:1 to the original toehold strand amount supplemented with additional MgCl_2_ and KCl to a final concentration of 2 mM and 50 mM, respectively, in a 30 µL reaction.

### Renaming and deleting

Toehold strands were mixed with renaming or deleting oligos at a ratio of 1:20 in a 25 µL reaction containing 2 mM MgCl_2_ and 50 mM KCl. The mixture was heated to 35 °C for 2 min, followed by a temperature drop at 1 °C/min to 14 °C. To delete the file, oligos were mixed with purified target file strands at a ratio of 1:20.

### Real-Time PCR (qPCR)

qPCR was performed in a 6 μL, 384-well plate format using SsoAdvanced Universal SYBR Green Supermix (BioRad, 1725270). The amplification conditions were 95 °C for 2 min and then 50 cycles of: 95 °C for 15 s, 51.5 °C for 20 s, and 60 °C for 20 s. Quantities were interpolated from the linear ranges of standard curves performed on the same qPCR plate.

### Theoretical thermodynamic calculations

To theoretically estimate the fraction of bound oligos with various overhang lengths and at different temperatures, we calculated the equilibrium constants at each condition:

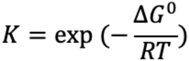

Where ΔG^0^ is the change in Gibbs Free Energy at standard conditions (25 °C, pH=7 in this case); R is the gas constant and T is the reaction temperature. The Gibbs Free Energy for each oligo was obtained using the Oligonucleotide Properties Calculator^33–35^. The equilibrium constant at each condition was equated to

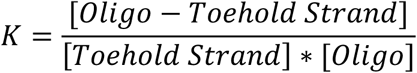

with:

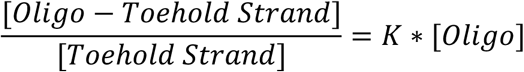

representing the fraction of accessed strands (strands separated out) to the total original amount of toehold strands. This amount, expressed as a percentage, is referred to as the access efficiency.

### Density and capacity calculation

Experimental work was performed using the oligos listed in Table S1. Simulation densities were measured by calculating the number of bytes in a 160 bp data payload with 5 codewords used for the strand index^11^, with the codeword length given as L:

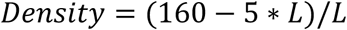

The size of the index is chosen to accommodate 10^9^ strands.

Capacity: For each density and corresponding number of oligos, system capacity is calculated assuming 10^9^ strands per file, which roughly corresponds to the number of strands that can be sequenced at a time in next generation sequencing. We further assume that each strand occurs 10 times in replicate.

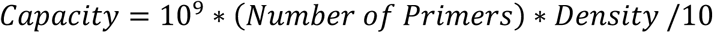

Note, these capacity calculations are based on the number of oligos found in our search (Figure S7), not the total number that may be available if we searched the entire space of all possible 20 bp oligos. Searching for more oligos will result in greater system capacity, but searching the entire space is intractable using our current approach.

## Acknowledgements

We acknowledge Kyle J. Tomek and Kevin Volkel for helpful discussions. We thank Prof. Nathan Crook for use of their Qubit fluorometer and reagents. This work was supported by the National Science Foundation (CNS-1650148 & CNS-1901324), a North Carolina State University Research and Innovation Seed Funding Award (#2018-2509), and a North Carolina Biotechnology Center Flash Grant to AJK and JT. KNL was supported by a Department of Education Graduate Assistance in Areas of Need fellowship.

## Author Contributions

KNL, JMT, and AJK conceived the study. KNL planned and performed the wetlab experiments with guidance from AJK. JMT planned and performed the simulations. KNL and AJK wrote the paper with input from all.

