## Supplementary Data for "Dynamic DNA-based information storage"

### SUPPLEMENTARY INFORMATION

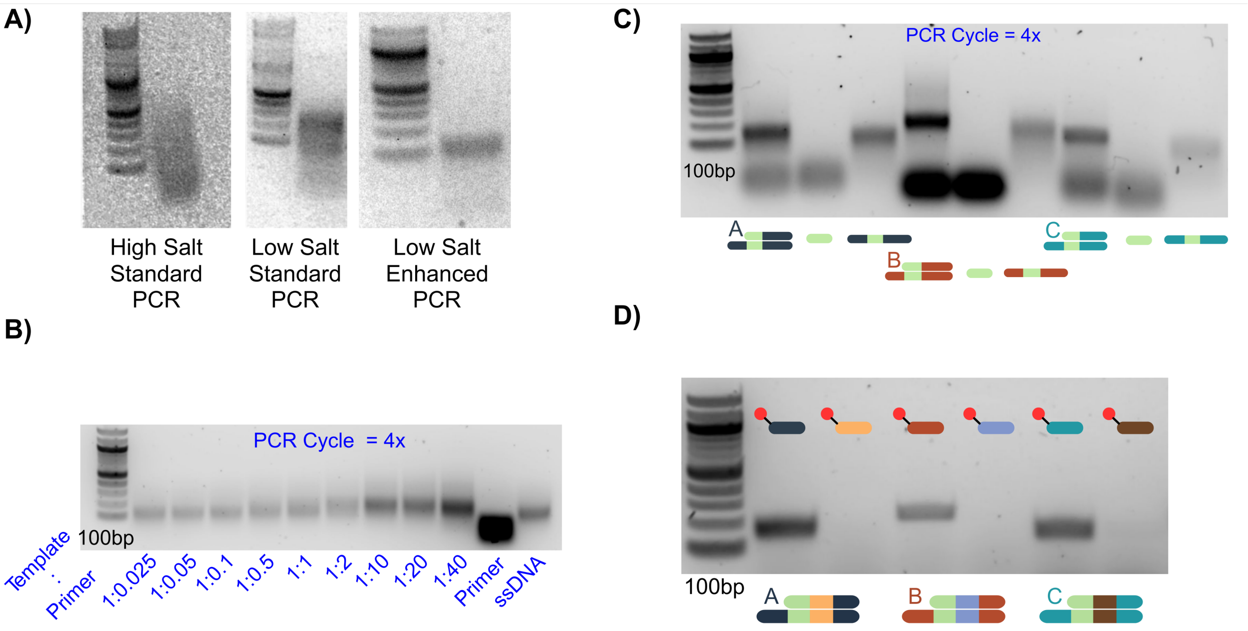


**Figure S1. Optimization of toehold generation through single primer extension.**

**A)** DNA gel electrophoresis. A low salt "enhanced" PCR buffer condition was identified that yielded a tight band corresponding to toehold strands. Other PCR buffers resulted in multiple byproducts (buffer details described in Methods). **B)** Increasing the amount of primer relative to ssDNA template in the primer extension process results in increasing amounts of toehold strands generated, as seen in a shift upwards in size from ssDNA. A slight downward shift at ratios of 1:40 suggest production of excess ssDNA. **C)** Toehold structures, primers, and ssDNA templates were run on a DNA gel to show their differences in band locations. **D)** File access oligos complementary to each toehold strand's address or to the middle of the toehold strand were mixed with samples after primer extension and then separated using magnetic beads. The absence of DNA when using oligos complementary to the middle of the strands indicate that the primer extension step was nearly complete in converting the ssDNA templates to toehold strands, thus blocking 'non-specific' file access.

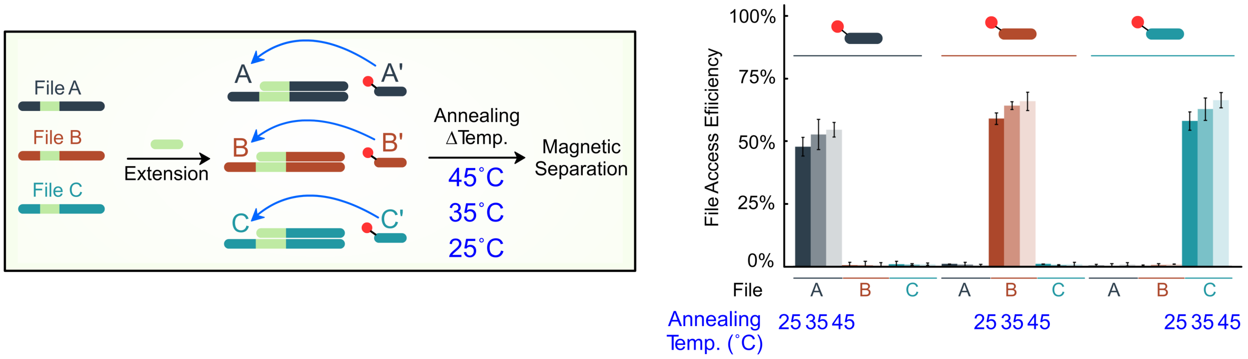


**Figure S2. File access temperature does not appreciably affect access efficiency from a 3-file database created through 'one-pot' primer extension.**

After the 3-file database was created, each file was accessed by its corresponding biotin-linked oligo at 25, 35, or 45 ˚C and separated using magnetic beads. The amounts of each file in the samples were quantified by qPCR. Each oligo accessed its file specifically, and the access/annealing temperature did not appreciably affect the file access efficiency, which was calculated as the amount of DNA accessed relative to its original quantity in the database. Error bars are standard deviations of three replicate file accesses/separations.

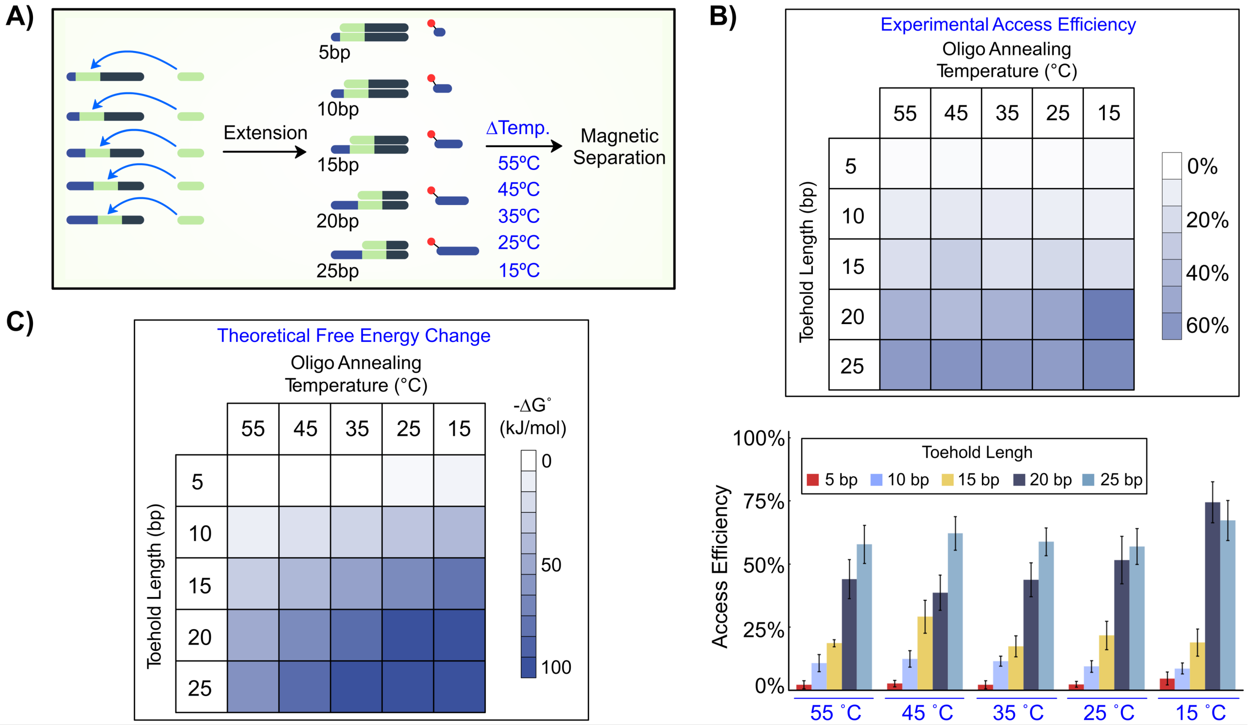


**Figure S3. Experimental and theoretical analyses identify the dependency of file access efficiency on oligo length and access temperature.**

**A)** 5 toehold lengths were generated by single primer extension and then evaluated for their file access efficiencies at 5 different temperatures. **B)** Experimental analysis of access efficiency displayed as both heatmap and bar graph. The access efficiency was calculated as the amount of file accessed relative to its starting total quantity as measured by qPCR. **C)** A theoretical analysis of the change in Gibbs free energy at different oligo/toehold lengths. Error bars are standard deviations of three replicate file accesses/separations.

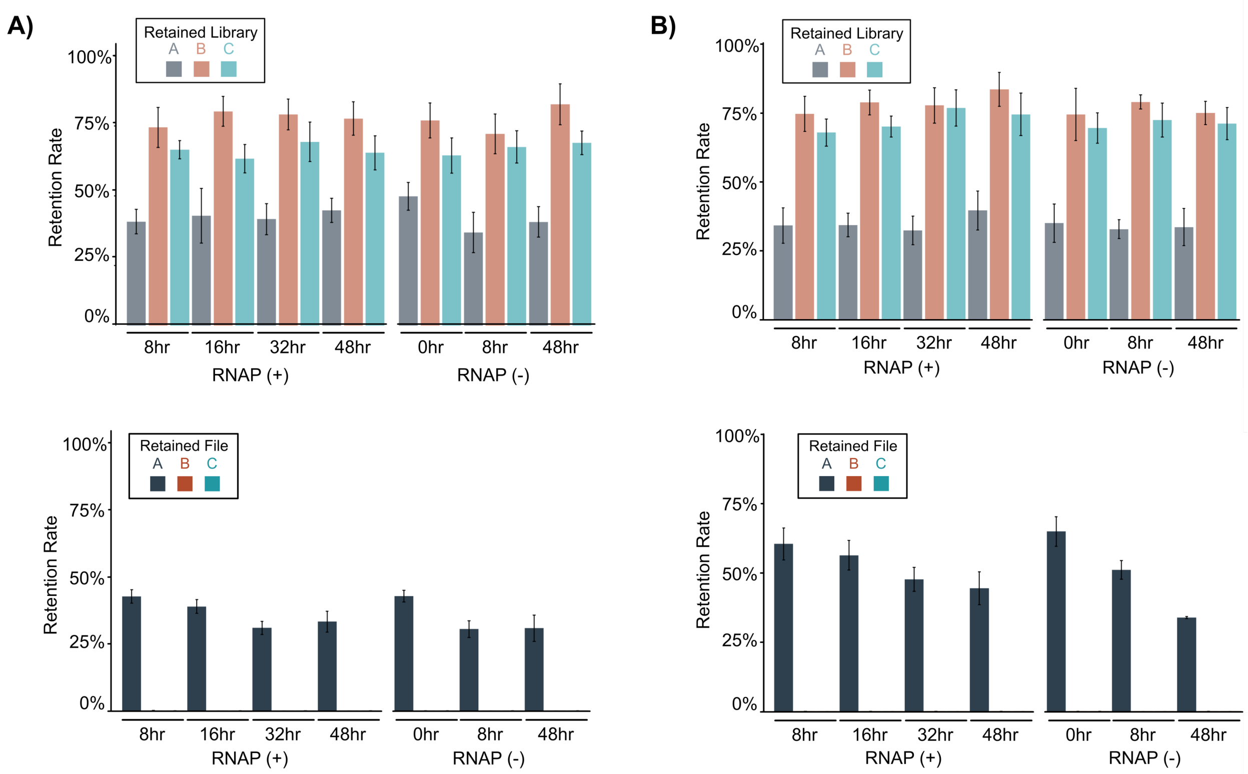


**Figure S4. IVT time but not the presence of RNA polymerase decreases the amount of Retained File. A, B)** The presence/absence of RNA polymerease and IVT time did not affect the retention rate of the Retained Library sample as these samples did not undergo IVT. The presence/absence of RNA polymerase did not affect the retention rate of the Retained File; however, the IVT time did. The decrease in retention rate of the retained file may be partially due to disrupted file-to-bead binding during the elevated temperature of the IVT step. **B)** A re-annealing step to 45 ˚C was able to rescue some of this loss. Retention rate is the amount of DNA recovered relative to the starting amount of DNA. Error bars are standard deviations of three replicate IVTs.

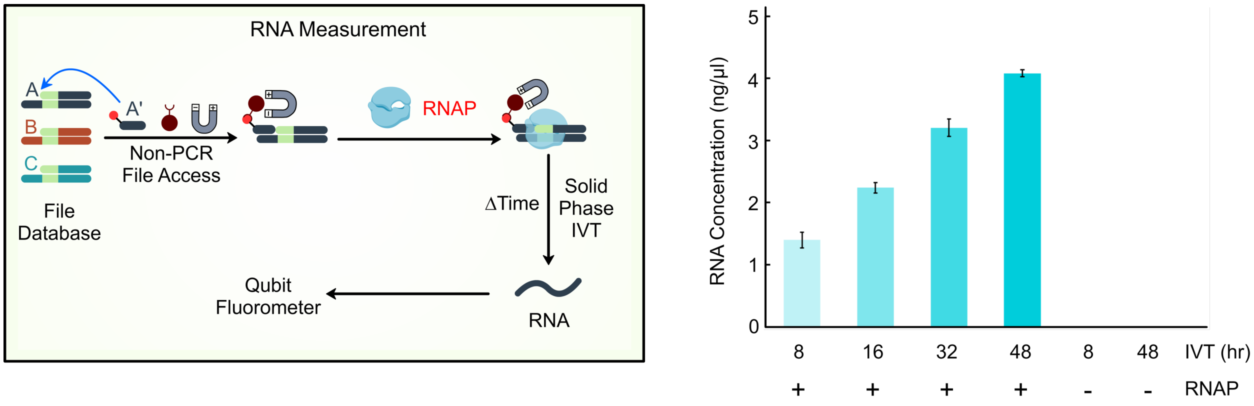


**Figure S5. Increasing IVT time increases the quantity of RNA produced.**

IVT was performed directly on File A strands bound to magnetic beads for different lengths of time, with or without RNA polymerase. The amount of RNA produced was measured directly by a Qubit fluorometer. Error bars are standard deviations of three replicate separated files and their corresponding IVTs.

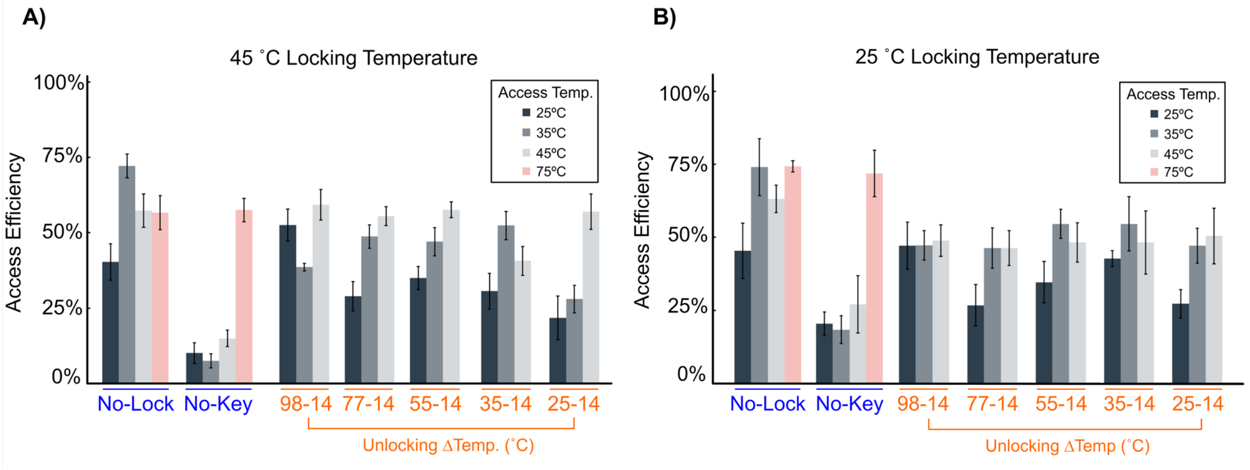


**Figure S6. Temperature influences the extent of locking.**

File A was accessed by DORIS without locking, or, following provision of a lock, was accessed with or without subsequent unlocking by a key. The lock was added at **A)** 45 °C or **B)** 25 °C and then cooled to 14 °C. The accessing oligo A' was added at different access temperatures of 25, 35, 45, or 75 ˚C for 2min, followed by a temperature drop of 1 ˚C/min to 25 °C. Access efficiency is the amount of file A recovered relative to its original quantity, as measured by qPCR. Error bars are standard deviations of three replicate file operations/accesses.

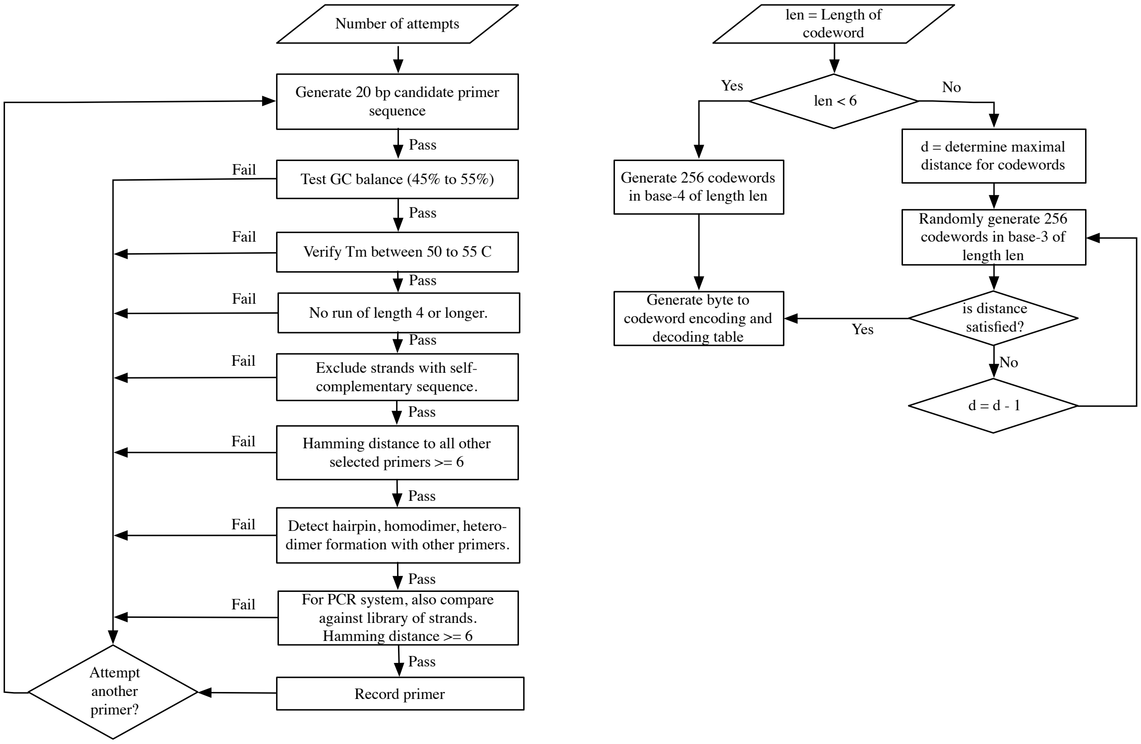


**Figure S7. Flowcharts for estimating total number of primers and for producing coding tables at varying density.**

**(Left)** Flowchart for estimating viability of a primer implemented as a Python program. The overall process was a loop that repeated over some number of attempts to find a primer. Primer sequences were generated at random according to a uniform distribution of A, C, G, and T. Then, each primer was evaluated against several criteria. For estimating Tm (melting temperature), hairpins, and other dimer formation, we used the Primer3 software[1]. We required that all primers were at least a Hamming distance of 6 apart, and we required that they were at least a Hamming distance of 6 away from the target library. We approximated that requirement by comparing each primer to 1 MB of data that was randomly generated and encoded in each run of the program. The encoding of the library was also an input to the process that could be varied to evaluate the impact of coding density on primer selection. **(Right)** Flowchart for producing encoding and decoding tables of varying length. The data payload of strands, used to validate primers in the flowchart on the left, were created by encoding each byte one at a time as a codeword. The codeword tables at various lengths were created through a common algorithm.

[1] Untergasser A, Cutcutache I, Koressaar T, et al. Primer3—new capabilities and interfaces. *Nucleic Acids Res*. 2012;40(15):e115-e115. doi:10.1093/nar/gks596

For length 4, all possible sequences are used, hence the creation is trivial. For length 5, we generate all possible sequences, and select 256 of them at random. For lengths of 6 or longer, the process is different. We generate codes in base-3 (ternary) and select 256 of them, one for each possible single byte value. To ensure the codewords were different enough from each other, we first attempt to generate codes of maximal Hamming distance according to the Singleton bound. If we do not succeed, we reduce the distance and try again. The codeword tables created by the algorithm were manually verified to have a distance of 2 or more for all lengths greater than 6. For length 6 and higher, after encoding an entire strand, we use a rotating encoding to ensure no repetitions in the strand, neither within nor across codewords.

**Table S1. Template and oligo design.**

| DNA Oligo | Sequence |
|---|---|
| ssDNA (File A) | CGTACGTACGTACGTCGACGGATGACAGCTCGCATCTACGAGCTCGAG ATGACACAGAGTATCGCATCTACGACACAGTCTCTCGCGAGCTAGAGA TGAGTGATCGAGCTCTGCTCGGCGCGCTATAGTGAGTCGTATTACGAG TGCAGAGCAGACTCAC |
| ssDNA (File A-2 for Truncated PCR) | CGTACGTACGTACGTCGACGGATGACAGCTCGCATCTACGAGCTCGAG ATGACACAGAGTATCGCATCGAGTGCAGAGCAGACTCACAGCTAGAG ATGAGTGATCGAGCTCTGCTCGGCGCG**C**TATAGTGAGTCGTATTACGA GTGCAGAGCAGACTCAC |
| ssDNA (File B) | CAGGTACGCAGTTAGCACTCCGTACGTACGTACGCAGCTAGCTCGATG AGTACTCTGCTCGATGAGTACTCTGCTCGACGAGATGAGACGAGTCTC TCGTAGACGAGAGCAGACTCAGTCATCGCGCTAGAGAGCATAGAGTC GTGATCTATGCTCAGCGCGCTATAGTGAGTCGTATTATCCGTAGTCATA TTGCCACG |
| ssDNA (File C) | GGGAGTAATCCCCTTGGCGGTCGCGGGGGACAGCGCGTACGTGCGTTT AAGCGGTGCTAGAGCTGTCTACGACCAGCGCGCGCTATAGTGAGTCGT ATTAGGATTCTCCAGGGCATCCGG |
| Extension Primer | TAATACGACTCACTATAGCGCGC |
| File Access Oligo A’ | GTGAGTCTGCTCTGCACTCG |
| File Access Oligo B’ | CGTGGCAATATGACTACGGA |
| File Access Oligo C’ | CCGGATGCCCTGGAGAATCC |
| PCR Oligo B | CTACGACACAGTCTCTCGCG |
| Reverse Oligo File A | CGTACGTACGTACGTCGACG |
| Reverse Oligo File B | CAGGTACGCAGTTAGCACTC |
| Reverse Oligo File C | GGGAGTAATCCCCTTGGCGGT |
| cDNA Forward Oligo | CGTACGTACGTACGTCGACG |
| cDNA Reverse Oligo | GAGCAGAGCTCGATCACTCA |
| File A Lock | CTCCATCAGAGTGATATGCCCAGCTTAGGTGAGTCTGCTCTGCACTCG |
| File A Key | CGAGTGCAGAGCAGACTCACCTAAGCTGGGCATATCACTCTGATGGAG |
| File A-> B Rename Oligo | TCCGTAGTCATATTGCCACGGTGAGTCTGCTCTGCACTCG |
| File A-> C Rename Oligo | GGATTCTCCAGGGCATCCGGGTGAGTCTGCTCTGCACTCG |
| File A Delete Oligo | GTGAGTCTGCTCTGCACTCG |
